# Propionylpromazine: Unveiling a Novel Inhibitory Agent Against *Mycobacterium tuberculosis*

**DOI:** 10.1101/2024.04.16.589721

**Authors:** U. Suresh, M. Punnam, C. Kartick, A. P. Sugunan

## Abstract

The emergence of multidrug-resistant *mycobacterium tuberculosis* presents a significant hurdle, as it shows resistance to existing standard therapies, underscoring the critical need for novel antibacterial agents. This study presents propionylpromazine as a promising compound with potent activity against *M. tuberculosis*. Our research indicates that propionylpromazine effectively curtails the proliferation of *M. tuberculosis* H37Rv. Importantly, propionylpromazine has shown the capability to impede the growth of *M. tuberculosis* within macrophages without causing detrimental effects. These findings endorse the potential of propionylpromazine to be further developed as a clinical drug for the effective treatment of *M. tuberculosis* infections.

## INTRODUCTION

Since 2014, Tuberculosis (TB), caused by *Mycobacterium tuberculosis*, has been a leading cause of death from a single infectious agent. The World Health Organization (WHO) reported that TB affected approximately 10 million people globally and resulted in at least 1.45 million deaths in 2018 (Chakaya et al., 2021). Notably, about 484,000 of these TB cases were resistant to rifampin, with 78% being classified as multidrug-resistant TB (MDR-TB). The current treatment for MDR-TB has a global success rate of only 56%, dropping to about 41% in China (Bagcchi, 2023). The latest WHO guidelines suggest a shorter treatment duration of 9–12 months for MDR-TB (Organization, 2013). However, the complexity and length of this treatment lead to poor patient compliance. This underscores the urgent need for the development of new drugs and treatment regimens that are shorter and simpler, to improve TB management. Recently, several new chemical entities and antimicrobial compounds have shown promise against the highly drug-resistant *M. tuberculosis*, including ganfeborole, DprE1 inhibitors, Q203, MmpL3 inhibitors, gwanakosides A, retinestatin, etc (Piton et al., n.d.; Li et al., 2016; Huynh et al., 2022, 2023; Nguyen et al., 2022).

Propionylpromazine, a variant of the phenothiazine class (Figure 1), serves as a calming and sleep-inducing agent in animal healthcare (Papich, 2016). This compound is a modified form of promazine, an antipsychotic medication, designed with a propionyl group to suit veterinary applications (Papich, 2016). Propionylpromazine hydrochloride has garnered attention for its possible antibacterial properties, particularly against tough, antibiotic-resistant bacteria such as *Acinetobacter baumannii* (Mohammed et al., 2020). This pathogen poses a significant health risk as it can cause severe infections that are notoriously challenging to treat because of its ability to withstand many conventional antibiotics (Mohammed et al., 2020). Research into PPZHCl’s effectiveness in inhibiting the growth of such resistant bacterial strains could lead to new avenues for treating infections that currently have limited therapeutic options. Nevertheless, the precise mechanisms of action are still undergoing investigation (Papich, 2016). There is no study about propionylpromazine against *M. tuberculosis*.

**Figure 1.**
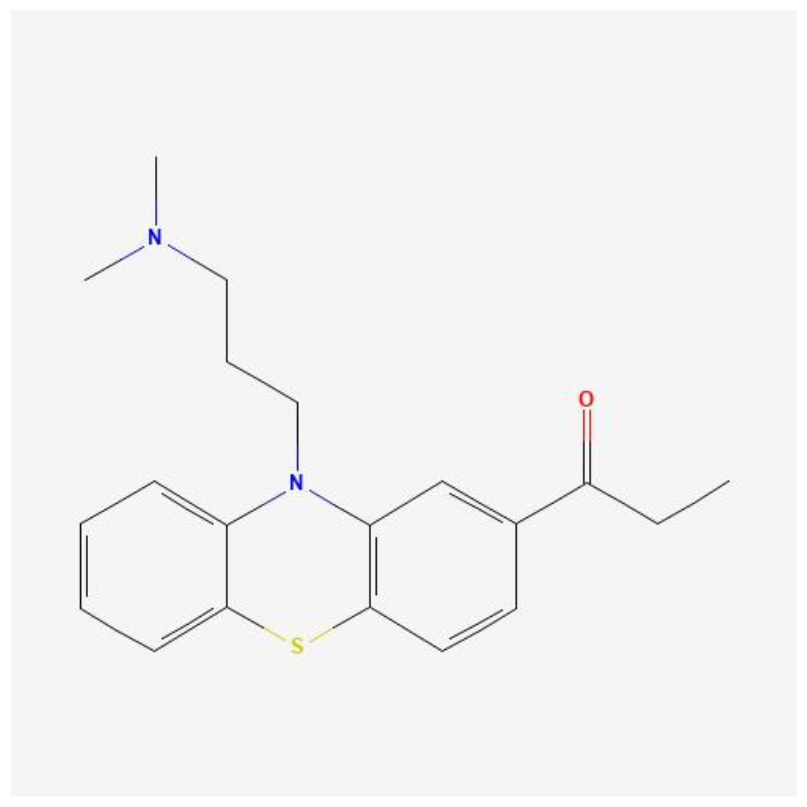
Chemical structure of propionylpromazine

In our findings, we present the discovery of a novel agent called propionylpromazine, which demonstrates remarkable efficacy against *M. tuberculosis* both in vitro and in infected murine macrophages.

## MATERIALS AND METHODS

### Bacterial strains / culture conditions / chemicals

*M. tuberculosis H37Rv* (ATCC 27294), stored at −80 °C, was carefully regrown in Middlebrook 7H9 broth. This nutrient-rich medium was enhanced with a 10% solution of oleic-acid-albumin-dextrose-catalase, along with 0.2% glycerol and 0.05% Tween 80, and was incubated at a temperature of 37 °C to promote bacterial growth. All drugs used in the study were bought from Meilun bio, Dalian, China.

### Resazurin Microtiter Assay (REMA)

The determination of MIC values was carried out using the resazurin microtiter assay (REMA). In this assay, bacterial stocks from the exponential-phase cultures were diluted to an optical density at 600 nm (OD600) of 0.01. Each well of a sterile, polystyrene 96-well cell culture plate was then filled with 50 μL of the bacterial culture, and 50 μL of serial 2-fold dilutions of the test compound solution were added to each well. A drug-free growth control was included on each plate. To prevent evaporation during incubation, 200 μL of sterile water was added to the outer perimeter wells. The plates were covered with self-adhesive membranes and incubated at 37°C for 5 days. Afterward, 40 μL of a 0.025% (wt/vol) resazurin solution was added to each well, and the plates were reincubated overnight. The fluorescence was measured using a SpectraMax M3 multimode microplate reader (Molecular Devices, Sunnyvale, CA, USA). A dose-response curve was constructed, and the concentrations required to inhibit bacterial growth by 50% (MIC50) were determined using GraphPad Prism software (version 6.05; San Diego, CA, USA).

### Isolation of BMDMs and intracellular bacterial replication assay

Bone marrow-derived macrophages (BMDMs) were obtained by flushing the femur and tibia of 6-week-old C57BL/6 mice. The BMDMs were cultured in Dulbecco’s modified Eagle’s medium (DMEM; Welgene, Gyeongsan-si, Gyeongsangbuk-do, South Korea) supplemented with 10% fetal bovine serum (FBS; Welgene), GlutaMax (35050-061; Gibco), and penicillin/streptomycin (15140-122; Gibco) at 37°C with 5% CO2. To differentiate the cells into mature BMDMs, they were exposed to recombinant murine macrophage colony-stimulating factor (M-CSF; JW-M003-0025, JW CreaGene) for a period of 5 days.

To test the antibacterial activity of propionylpromazine against intracellular bacteria, the BMDMs were infected with *M. tuberculosis* subsp. *abscessus* ATCC 19977 at a multiplicity of infection of 1 in 96-well plates for a duration of 3 hours. After the infection, extracellular bacteria were eliminated by washing three times using phosphate-buffered saline (PBS; Gibco) and treated with serially diluted compounds in 96-well plates for a period of 3 days. To release the intracellular bacteria, the cells were lysed using 1% SDS (151-21-3; Generay Biotechnology). The lysates were subsequently 10-fold serially diluted with PBS, and each dilution was plated on 7H10-OADC supplemented with 50 μg/mL kanamycin. The plates were incubated for at least 3 days at 37°C, after which the bacterial colonies were counted.

## RESULTS

### Propionylpromazine Exhibits Potent Activity against M. tuberculosis H37Rv

The effectiveness of propionylpromazine against *M. tuberculosis H37Rv* was evaluated by identifying the minimum concentration needed to inhibit 50% of bacterial growth. Bedaquiline served as a benchmark for comparison. Following a 5-day incubation period with these compounds, a reduction in fluorescence signaled a decrease in bacterial viability. The results, depicted in Figure 2, showed that *M. tuberculosis H37Rv* was vulnerable to both propionylpromazine, with an MIC50 of 1.4 μM and an MIC90 of 3.2 μM, and bedaquiline, which exhibited a comparable MIC50. These findings suggest that propionylpromazine is a promising candidate for treating *M. tuberculosis* infections.

**Figure 2.**
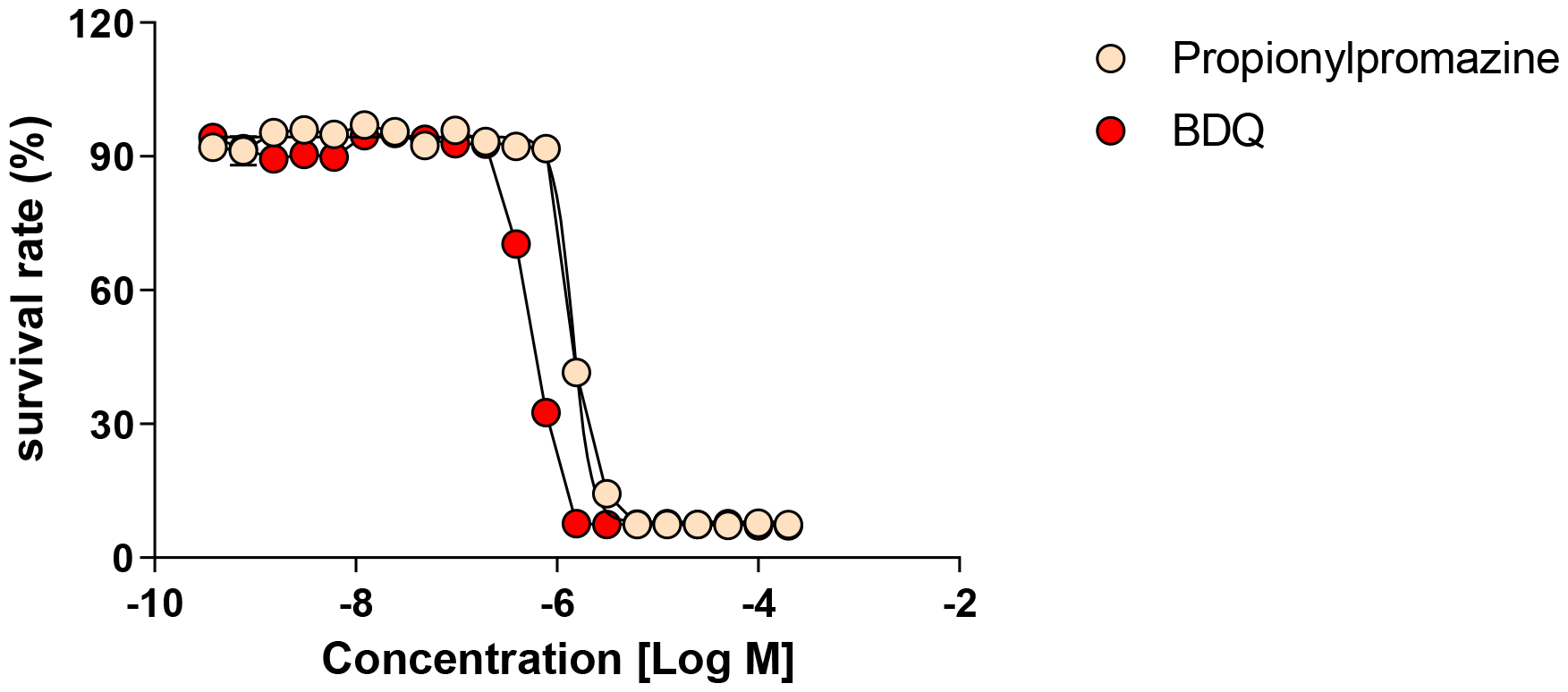
In vitro activity of propionylpromazine against *M. tuberculosis H37Rv*

**Figure 3.**
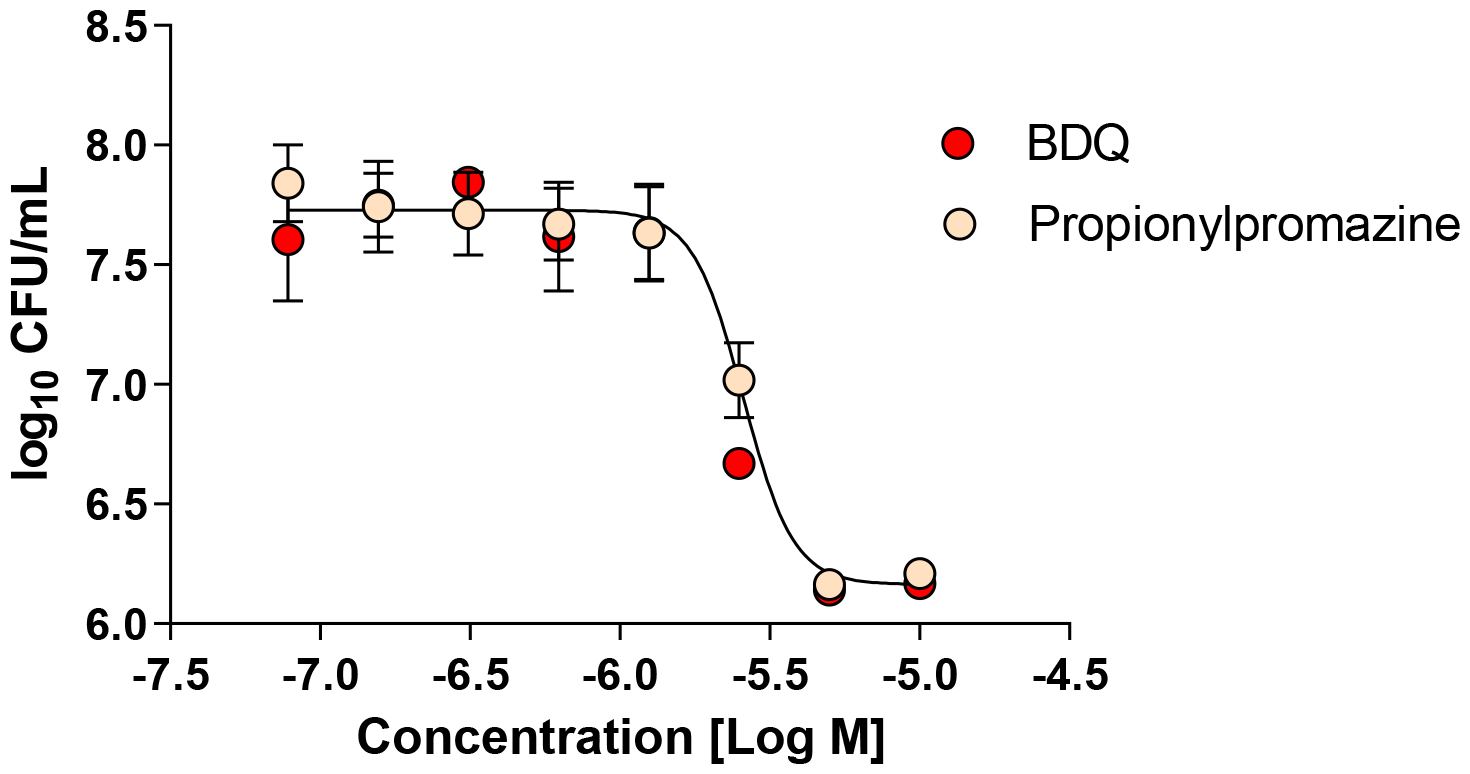
Intracellular activity of propionylpromazine against *M. abscessus*

### Propionylpromazine activity against intracellular replicating M. tuberculosis H37Rv

To evaluate propionylpromazine’s ability to inhibit M. tuberculosis replication in host cells, researchers infected mouse bone marrow-derived macrophages (mBMDMs) with the bacterium. Bedaquiline served as the positive control, while dimethyl sulfoxide (DMSO) was the negative control. The traditional colony count method was used to measure the number of viable M. tuberculosis after lysing the mBMDMs treated with propionylpromazine and clarithromycin. The study’s results indicated a notable reduction in intracellular M. tuberculosis numbers due to propionylpromazine treatment, with a potency similar to that of bedaquiline, as reflected by IC50 values of 2.5 μM for propionylpromazine and 2.2 μM for bedaquiline. These outcomes suggest that propionylpromazine is effective against M. tuberculosis within host cells.

## DISCUSSION

Tuberculosis (TB) remains a significant global health challenge, with the World Health Organization (WHO) reporting 10 million cases and 1.45 million deaths in 2018 (Bagcchi, 2023). A concerning aspect is the prevalence of rifampin-resistant TB cases, with 78% being multidrug-resistant (MDR-TB) (Chakaya et al., 2021). The treatment success rates for MDR-TB are low, particularly in regions like China. Recognizing the issues with the lengthy and complex treatment regimens that contribute to poor patient adherence, the WHO now recommends a shorter 9–12 month treatment course for MDR-TB (Chakaya et al., 2021). Despite this, the need for new, more effective, and simpler treatments is critical. Promising research into new chemical entities and antimicrobial compounds, such as ganfeborole, DprE1 inhibitors, Q203, MmpL3 inhibitors, gwanakosides A, and retinestatin, offers hope for improved management of drug-resistant TB strains (Piton et al., n.d.; Li et al., 2016; Huynh et al., 2022, 2023; Nguyen et al., 2022).

Propionylpromazine, a phenothiazine derivative, is utilized in veterinary medicine for its sedative effects (Papich, 2016). It is a modified version of the antipsychotic drug promazine, tailored with a propionyl group for animal treatment (Papich, 2016). The hydrochloride form of propionylpromazine has sparked interest due to its potential to combat antibiotic-resistant bacteria, such as Acinetobacter baumannii, which is notorious for causing hard-to-treat infections (Mohammed et al., 2020). Ongoing research into its antibacterial efficacy could pave the way for new treatments for infections with limited current options. However, the specific mechanisms by which it works are still being studied, and there are no studies to date on its effects against *M. tuberculosis*. In our investigation, we evaluated propionylpromazine’s impact on *M. tuberculosis* and found it to be consistently effective. Our initial tests on the in vitro response of *M. tuberculosis* to propionylpromazine showed a substantial decrease in bacterial survival, with its efficacy on par with bedaquiline. These outcomes underscore propionylpromazine’s promise as a potential therapeutic for *M. tuberculosis* infections.

The research focused on the ability of propionylpromazine to act as an intracellular antimicrobial against *M. tuberculosis* within murine bone marrow-derived macrophages (mBMDMs). *M. tuberculosis* typically avoids immune detection by replicating inside these immune cells. The study found that propionylpromazine significantly reduced the number of *M. tuberculosis* cells inside mBMDMs, showing effectiveness similar to bedaquiline. This suggests that propionylpromazine has potential as an intracellular antimicrobial agent. The promising results support further investigation of propionylpromazine as a potential treatment for *M. tuberculosis* infections, warranting additional research and clinical testing to confirm its therapeutic value and safety profile.

## Supporting information

raw data

## AUTHOR CONTRIBUTIONS

APS concptualised the research work. US, MP, and CK searched and gathered the previous studies. APS wrote the manuscript. US, MP, and CK critically reviewed the manuscript. RB edited the reviewed manuscript. APS critically evaluated and revised the manuscript and supervised the whole project.

## ACKNOWLEDGMENTS

The authors would like to acknowledge the Indian Council of Medical Research (ICMR), New Delhi, for their financial support in conducting this study. Their support has been instrumental in enabling the research and contributing to the findings presented in this work.

