## Supplementary material for "Propionylpromazine: Unveiling a Novel Inhibitory Agent Against *Mycobacterium tuberculosis*": raw data: raw data.pptx

### Slide 1
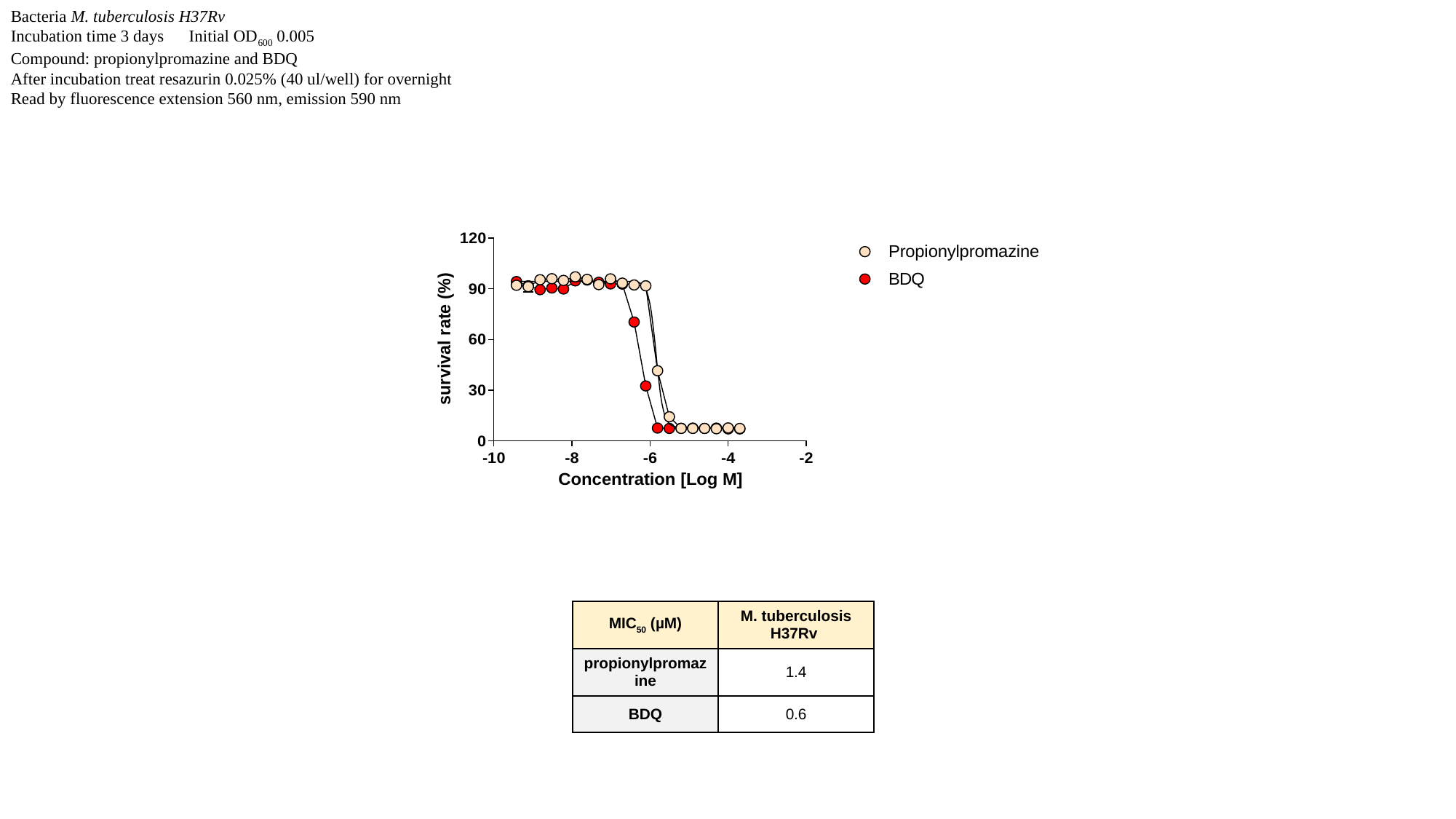

Bacteria M. tuberculosis H37Rv
Incubation time 3 days Initial OD_600_ 0.005
Compound: propionylpromazine and BDQ
After incubation treat resazurin 0.025% (40 ul/well) for overnight
Read by fluorescence extension 560 nm, emission 590 nm
| MIC_50_ (µM) | M. tuberculosis H37Rv |
| --- | --- |
| propionylpromazine | 1.4 |
| BDQ | 0.6 |

### Slide 2
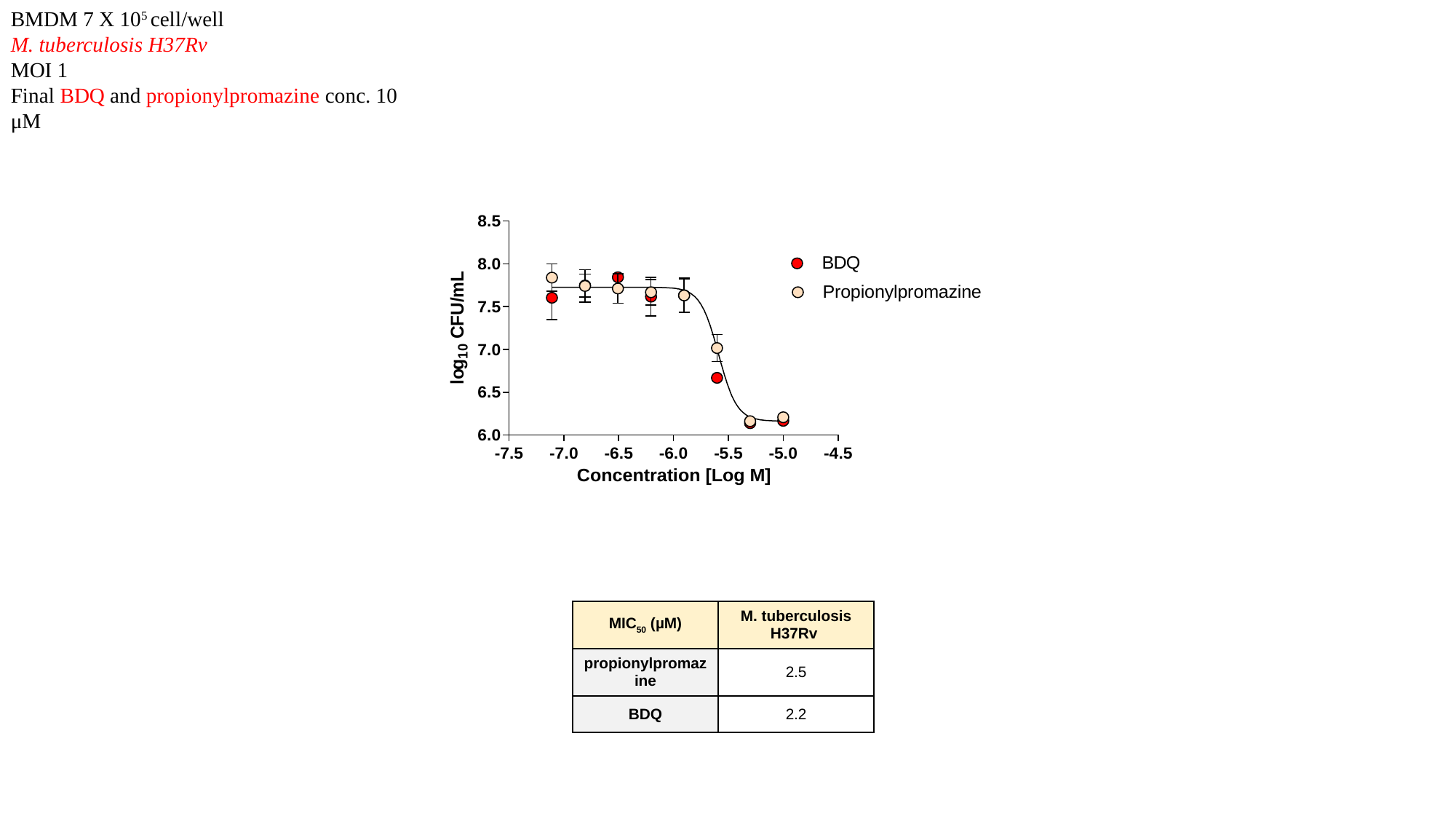

BMDM 7 X 10^5^ cell/well
M. tuberculosis H37Rv
MOI 1
Final BDQ and propionylpromazine conc. 10 μM
| MIC_50_ (µM) | M. tuberculosis H37Rv |
| --- | --- |
| propionylpromazine | 2.5 |
| BDQ | 2.2 |
